# ARPEGGIO: Automated Reproducible Polyploid EpiGenetic GuIdance workflOw

**DOI:** 10.1101/2020.07.16.206193

**Authors:** Stefan Milosavljevic, Tony Kuo, Samuele Decarli, Lucas Mohn, Jun Sese, Kentaro K. Shimizu, Rie Shimizu-Inatsugi, Mark D. Robinson

## Abstract

Whole genome duplication (WGD) events are common in the evolutionary history of many living organisms. For decades, researchers have been trying to understand the genetic and epigenetic impact of WGD and its underlying molecular mechanisms. Particular attention was given to allopolyploid study systems, species resulting from an hybridization event accompanied by WGD. Investigating the mechanisms behind the survival of a newly formed allopolyploid highlighted the key role of DNA methylation. With the improvement of high-throughput methods, such as whole genome bisulfite sequencing (WGBS), an opportunity opened to further understand the role of DNA methylation at a larger scale and higher resolution. However, only a few studies have applied WGBS to allopolyploids, which might be due to lack of genomic resources combined with a burdensome data analysis process. To overcome these problems, we developed the Automated Reproducible Polyploid EpiGenetic GuIdance workflOw (ARPEGGIO): the first workflow for the analysis of epigenetic data in polyploids. This workflow analyzes WGBS data from allopolyploid species via the genome assemblies of the allopolyploid’s parent species. ARPEGGIO utilizes an updated read classification algorithm (EAGLE-RC), to tackle the challenge of sequence similarity amongst parental genomes. ARPEGGIO offers automation, but more importantly, a complete set of analyses including spot checks starting from raw WGBS data: quality checks, trimming, alignment, methylation extraction, statistical analyses and downstream analyses. A full run of ARPEGGIO outputs a list of genes showing differential methylation. ARPEGGIO’s design focuses on ease of use and reproducibility. ARPEGGIO was made simple to set up, run and interpret, and its implementation includes both package management and containerization. Here we discuss all the steps, challenges and implementation strategies; example datasets are provided to show how to use ARPEGGIO. In addition, we also test EAGLE-RC with publicly available datasets given a ground truth, and we show that EAGLE-RC decreases the error rate by 3 to 4 times compared to standard approaches. The goal of ARPEGGIO is to promote, support and improve polyploid research with a reproducible and automated set of analyses in a convenient implementation.

## Background

Polyploidy, also known as whole genome duplication (WGD), is a process leading to the formation of an organism with more than two sets of chromosomes. There are two types of polyploidy: autopolyploidy, the doubling of an entire genome in a single species, and allopolyploidy, the hybridization of two different species followed by whole genome duplication (1). Both of these processes influenced the evolutionary history of many living organisms such as nematodes, arthropods, chordates, fungi, oomycetes and plants (1–3). Of all these lineages, the most extensive research on polyploidy has been done on land plants (1–8), where about 35% of all species were estimated to be recent polyploids (7,8) and at least one ancient WGD was inferred in the ancestry of every lineage (3).

To understand the successful prevalence of WGD and the underlying mechanisms, particular attention was given to early stages of polyploidy in allopolyploids (4,9–11). Among several observed genomic and epigenomic changes (4,10,12), DNA methylation was shown to play an important role to ensure the survival of a newly formed allopolyploid (13–19). A well-studied example comes from Madlung and colleagues (13) in which they chemically treated synthetic *Arabidopsis suecica* allotetraploids to remove DNA methylation over the whole genome. With this treatment, they observed many phenotypic disorders such as abnormal branching or homeotic abnormalities in flowers, mostly leading to sterility. These abnormalities were not observed when treating the parent species or the natural allopolyploid, highlighting the importance of DNA methylation in the first generations after allopolyploidization. Follow-up studies focused on the epigenetic regulation in other resynthesized allopolyploid species with varying outcomes. In allopolyploid wheat, *Tragopogon, Spartina* and rice, DNA methylation changes indicated gene repression favoring one parental genome over the other (15–20). This was not the case in *Arabidopsis*, where similar DNA methylation and expression changes were observed on both parental genomes (21). In *Brassica*, both previously mentioned outcomes were reported (15,22), while in cotton no changes were found (23). All these studies proposed different mechanisms to clarify the role of methylation and its short and long term evolutionary impact, but the discussion remains open (4). One reason that might complicate the grounds of such discussion, is the variety of tools and methods used to analyze DNA methylation data. To better control discrepancies between findings caused by methodological differences, a standardized set of tools would be ideal.

Despite the potential significance of DNA methylation in allopolyploid evolution, many of the previously mentioned findings were limited by low-throughput methods. These methods, such as methylation-sensitive amplified length polymorphisms (MSAP), were unable to capture changes at a whole genome level (24). With advances in technology, new high-throughput methods such as whole genome bisulfite sequencing (WGBS) are able to obtain methylation information at individual nucleotides over the whole genome (25).

At the whole genome level, DNA methylation is separated into three different sequence contexts: CG, CHG and CHH (where H = A, T or C). Each context is regulated by different families of enzymes and depending on the species, some contexts might be more important than others (26). For example, in mammals, methylation occurs mainly in CG context, while in plants it occurs in all three contexts (26).

Although WGBS is considered to be the gold standard in whole-genome DNA methylation studies (24,27), research on allopolyploid species using WGBS is limited, with most of the studies coming from crop study systems (28–30). On the one hand, these systems have excellent genomic resources to provide valuable insights, while on the other, it is unclear whether these insights can be extended to wild organisms in nature given their artificial selection (4).

In other polyploid study systems, two major challenges prevent the use of WGBS: limited genomic resources (i.e. genome assemblies) and a laborious data analysis process. The number of plant genome assemblies has been increasing exponentially in the last years (31), but polyploid genome assemblies are still an intensive, complex and expensive task (32,33), preventing the development of genetic and epigenetic studies using polyploids. For allopolyploids, this obstacle can be avoided by using the genome assemblies of the two (known) parent species (34), usually diploid.

Besides limited genomic resources, another challenge in WGBS comes from a laborious and complex data analysis process (35–37). In standard WGBS data analysis pipelines, complexities related to polyploids are often not taken into account. For example when mapping reads originating from an allopolyploid, high sequence similarity between parents can be challenging for read mapping algorithms (38,39) and the outcome can have strong bias, especially when the quality of the assemblies is asymmetric (40). To tackle this problem, several methods were developed to improve the categorization of allopolyploids’ reads to the correct parental genome. HomeoRoq (41) and PolyDog (40) take into account alignment quality from both parental genomes to assign reads, while PolyCat (42) and EAGLE-RC (34) also use explicit genotype differences between parent genomes to classify reads. EAGLE-RC outperformed HomeoRoq in estimating homeolog expression with data from tetraploid *Arabidopsis* and hexaploid wheat (34). When comparing EAGLE-RC and PolyCat using *Gossypium* RNA-seq data, both tools outperformed other pipelines and had similar performance (43). Among all the tools, only PolyCat supports bisulfite-treated WGBS data, but only with available variant information (i.e. SNPs) between subgenomes, which represents an additional obstacle for most allopolyploid systems (44).

To promote and support allopolyploid DNA methylation research, we developed the Automated Reproducible Polyploid EpiGenetic GuIdance workflOw (ARPEGGIO). ARPEGGIO is a specialized workflow to process raw WGBS data utilizing the assemblies of the allopolyploid’s parent species (hereafter referred to as progenitors) or independently phased subgenomes of an allopolyploid. ARPEGGIO includes all the steps from raw WGBS data to a list of genes showing differential methylation: conversion check, quality check, trimming, alignment, read classification, methylation extraction, statistical analysis and downstream analysis. More details about the prerequisites, setup, tools and outputs are discussed in the implementation section.

To handle sequence similarity between two genomes, ARPEGGIO exploits an updated version of EAGLE-RC that supports bisulfite-treated reads and does not require variant information between subgenomes. This version of EAGLE-RC was evaluated using three WGBS datasets, and showed better performance compared to a genome concatenation approach.

ARPEGGIO’s implementation combines the Snakemake workflow management system (45) with the Conda package manager (46) and Singularity containers (47) to ensure both ease of use and reproducibility. For ease of use, a centralized configuration file controls all parameters related to ARPEGGIO and through Conda, all the tools required by the workflow are automatically installed.

## Implementation

### Design, concepts and challenges

ARPEGGIO’s design had three main objectives, each dealing with different aspects and challenges of the workflow: allopolyploid support, ease of use and reproducibility. These aspects will be discussed at high-level here and more details about their implementation can be found in the following sections.

To support allopolyploids, ARPEGGIO first needed to allow for different experimental designs (i.e. sample comparisons). For allopolyploids without a genome assembly, but progenitor assemblies available, there are two possible comparisons: allopolyploid against progenitors or allopolyploid against allopolyploid (Fig. 1a,b). The former compares the two allopolyploid’s subgenomes to the progenitors, while the latter compares directly the two subgenomes in different experimental conditions. An additional third comparison allows two groups of individuals from a species with an available (phased) genome assembly (Fig. 1c), regardless of the ploidy level. After choosing a comparison, the next allopolyploid-specific step is read classification.

**Figure 1:**
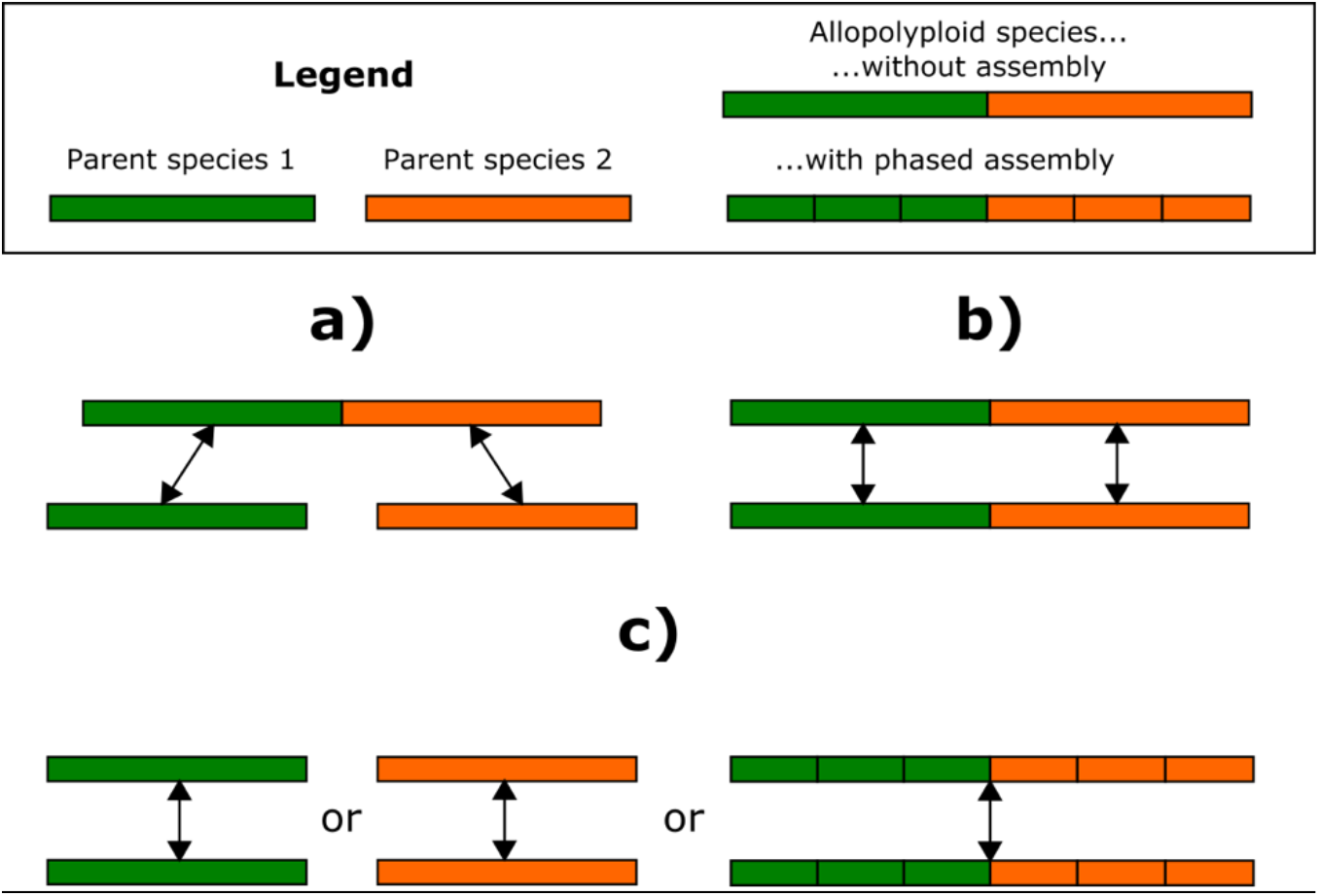
schematic view of the experimental designs supported by ARPEGGIO. There are 3 possible comparisons: a) polyploid species without assembly against its progenitors, b) same polyploid without assembly in two different experimental conditions and c) diploid species or polyploid species with an available phased assembly in two different experimental conditions. All comparisons are about whole genome DNA methylation patterns.

To analyze allopolyploid data with progenitor assemblies, we run two separate workflows in parallel, one per progenitor (Fig. 2). The separation occurs at the alignment and deduplication step, where two separate alignments are performed for the same allopolyploid data, one for each progenitor. With each allopolyploid read being mapped twice, a read classification algorithm must choose one of the two progenitors; tor the classification, ARPEGGIO uses EAGLE-RC. In short, EAGLE-RC applies a probabilistic method that compares the two mappings for each read and classifies its progenitor origin or deems it ambiguous (equal probabilities for both progenitors sides). Two parameters were added to EAGLE-RC to deal with bisulfite data from allopolyploids. The first is called “no genotype information” (NGI) and allows EAGLE-RC to be used with no information about variants in the genome. This mode is especially useful to reduce prerequisites for using ARPEGGIO. The second parameter is called “bisulfite” (BS) and it causes bisulfite treatment to be taken into account when a bisulfite-treated read is mapped to a genome. This parameter considers C-T as a match (forward strand), G-A as a match (reverse strand) or both.

**Figure 2:**
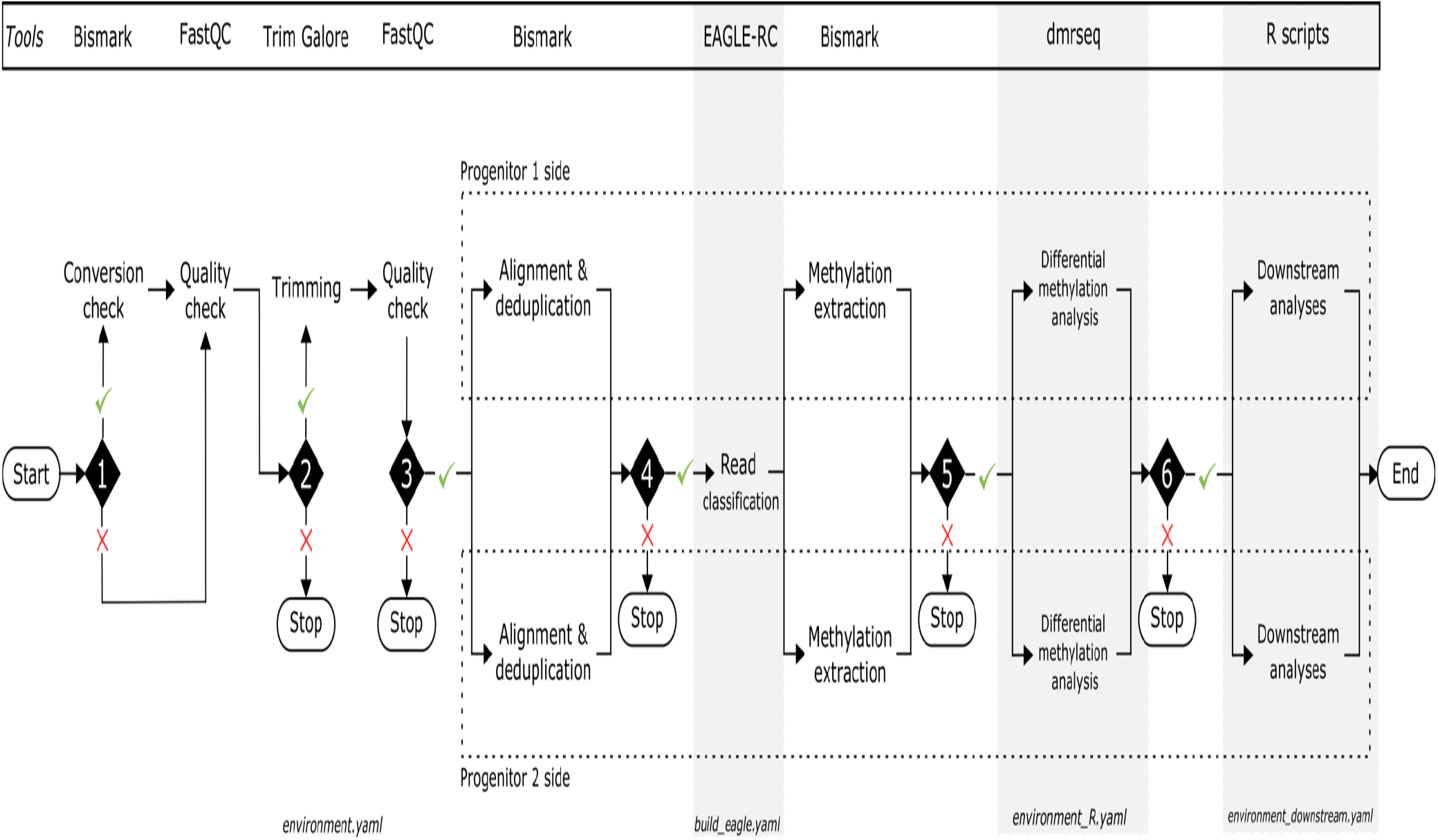
schematic overview of ARPEGGIO’s structure. All the shapes follow the flowchart standardized symbols (87). Ovals show the beginning and ends of the workflow. Diamonds represent conditional rules in ARPEGGIO’s configuration file and those rules make ARPEGGIO more adaptable to the needs of the user. Each conditional rule can be set to “true” (tick) or “false” (cross). Besides the first conditional rule, all other rules stop the workflow at the given point when set to “false”. The different grey backgrounds and the white background represent different Conda environments used by ARPEGGIO to carry out different steps of the analyses. In the scheme, the background of each step represents the environment that the step is part of. The bottom of each background shows the name of the file used to create the environment. At the top all the tools used by ARPEGGIO are shown and vertically aligned their corresponding step in the workflow. From the alignment and deduplication step, ARPEGGIO executes two workflows in parallel for each progenitor side, both highlighted by the dashed areas.

Both experimental design and EAGLE-RC’s inclusion had a major impact on ARPEGGIO’s structure and implementation, but other important aspects were also taken into account. For example, allopolyploids can be found in different lineages such as plants and mammals, meaning that different approaches should be considered for conversion efficiency checks and the selection of methylation contexts.

Once the general design of ARPEGGIO was established, the next challenge was to make the workflow easy to set up, run and interpret. ARPEGGIO requires the users to install the Conda package management system (46), then Snakemake (45) via Conda and, optionally, Singularity (48). No other tools need to be installed as ARPEGGIO will take care of automatically installing what is needed. To prepare ARPEGGIO for a new dataset, input files have to be prepared and ARPEGGIO’s settings have to be defined. Input files include raw data in FASTQ format and the progenitors’ reference genome assemblies. To run downstream analyses, annotation files for both assemblies are also required. ARPEGGIO’s settings are defined with a configuration file and a metadata file. The configuration file has different sections, each including parameters that define how ARPEGGIO will be run, while the metadata file contains information about samples such as filename, sequencing strategy, origin (allopolyploid or progenitor) and experimental condition (if present). A small dataset with its own configuration and metadata file are provided in ARPEGGIO’s repository as an example. To run ARPEGGIO, only one command is needed and its main options are related to reproducibility (discussed below) and parallelization (i.e. multiple core usage). After ARPEGGIO is successfully run, the number of files in the output folder can be significant. For this reason, a map of the output is available in ARPEGGIO’s user documentation: this map shows the general output structure with all the main folders and their contents. For each folder, there’s a section describing the folder itself, sub-folders and all the files included in it.

Another key goal of ARPEGGIO was to ensure reproducibility. Considering the variety of tools and the amount of steps in the workflow, by letting users (or Conda) define the version of each tool, the outcome could be variable and lead to future reproducibility problems. To overcome this, we fixed all the versions of the tools and we combined ARPEGGIO with Conda and Singularity containers. The user can choose to use either only Conda or Conda and Singularity together. The main difference between the two modes lies on potential issues between the user’s system and Conda. When these issues happen, Singularity offers a containerized run of Conda. Both these options can be specified with one or two parameters respectively when running ARPEGGIO. Aside from tool version differences, which we addressed above, the configuration file specifies all parameters that were used in a workflow run. Associating results to a specific set of parameters further aids reproducibility. The configuration file may also be shared to other researchers aiming to reanalyze a given dataset.

### Workflow overview

ARPEGGIO includes eight processes: conversion check, quality checks, trimming, alignment and deduplication, read classification, methylation extraction, differential methylation analysis and downstream analyses (Fig. 2). These processes are divided into six steps, each represented by a black diamond in Fig. 2. Step 1 includes conversion check, a quality check specific to WGBS data, where reads are aligned to an unmethylated control genome (usually plastid genome for plants and lambda genome for others) to assess the efficiency of the bisulfite conversion; the lower the mapping rate, the better the conversion (27). This process is executed by Bismark (49). The conversion check is followed by quality checks and trimming (step 1 and 2), executed by FastQC (50) and Trim Galore (51), respectively. Both processes are common procedures to assess read quality and remove noise. Step 3 performs read alignment to a reference genome, followed by deduplication, which removes duplicated reads. Both of these are carried out by the Bismark suite (49). From this point of the workflow allopolyploid data is separated into two parallel workflows: one per progenitor side. These workflows intersect in the next, allopolyploid-specific read classification step (step 4), executed by the updated version of EAGLE-RC (34). Here, EAGLE-RC will classify allopolyploid reads after comparing the read alignment on each progenitor’s side. After read classification (from step 5 on), the two workflows are independent, but execute the same steps. During methylation extraction via Bismark, methylation information is extracted for each cytosine from classified reads to produce a methylation count table. This table is used for differential methylation analyses (step 5), performed by the R/Bioconductor package `dmrseq` (52), to output a list of tested differentially methylated regions (DMRs). Finally, downstream analyses (step 6) consist of a series of R scripts for computing overlaps between statistically significant DMRs and annotated gene regions provided by the user (if available). More specifically, by default ARPEGGIO uses q-value < 0.05 to define a significant DMR. With this cutoff, ARPEGGIO looks for overlaps of at least 1 base pair between significant regions and gene regions based on the annotations. Before ARPEGGIO finishes a run, all reports (conversion check, quality checks, trimming, alignment, deduplication and methylation extraction) are combined into one interactive HTML report with MultiQC (53).

Each part in ARPEGGIO is optional and the user can specify which parts of the workflow to execute in the configuration file. It must be noted that skipping some parts will stop the workflow at a specific step (Fig. 2). Assuming that all prerequisites are met, ARPEGGIO goes from raw sequencing data to a list of genes showing differential methylation. Some useful intermediate outputs are also produced: an interactive HTML report merging all quality, alignment and methylation reports and an Rdata file with the output from the dmrseq analysis, which can be used to visualize DMRs or for other custom analyses.

### Implementation details

ARPEGGIO is written in Snakemake, a Python based language for workflow development (45). With Snakemake, a workflow is broken down into a series of rules. One rule can be seen as one step in the workflow with a defined input and output. Rules are related to each other based on their input and output files. Once all the rules are set, to run a Snakemake workflow, a target file (or multiple) needs to be requested. Snakemake will automatically build the workflow to obtain the target file based on the input/output relationships between rules (dependencies). If the relationships are successfully established, the workflow will be run. To illustrate these principles, an example with ARPEGGIO’s rules is given in Additional File 1. This figure shows all the input/output relationships between rules when running ARPEGGIO with single-end data, comparing an allopolyploid to its progenitor species (default experimental design).

In addition to the core features of Snakemake, ARPEGGIO takes advantage of the integrated Conda package management system (46). Conda creates environments containing a specific set of software and users can switch between different environments depending on the software package(s) they need. An environment can be created in several ways. ARPEGGIO creates environments through YAML files, specifying all the packages to be included and the channels from which the packages are searched. The integration of Conda in Snakemake allows rules to be run within a specific environment and during the execution of a workflow, Snakemake takes care of switching between environments if different rules require different environments. From a user perspective, once Conda and Snakemake are installed, ARPEGGIO will take care of installing all the tools needed for the analyses, running them and switching automatically between environments when needed (Fig. 2).

Making the workflow specific for allopolyploids presented major challenges with both Snakemake and Conda. Snakemake rules in ARPEGGIO had to be structured to allow for any combination between sequencing strategies and experimental designs. This meant combining rules for six workflows in one: three experimental designs, each with two sequencing strategies. In addition, since EAGLE-RC could not be installed as a Conda package, a Conda environment with a specific set of rules was created to take care of downloading, extracting and installing EAGLE-RC.

In practice, any user can take advantage of all the Conda and Snakemake features discussed above with a central configuration file. Here, we will discuss the first three sections of this file, that consist of parameters concerning the workflow as a whole: general parameters, conditional rules and experimental designs. All the other sections in the configuration file are related to tool-specific parameters for each of the main steps in ARPEGGIO. More details about these parameters can be found in ARPEGGIO’s user documentation. General parameters include the location of the output folder, the location of the metadata file and a parameter to define the sequencing strategy. Conditional rules are shown as black diamonds on Fig. 2. Those rules are set to “True” or “False” to define which parts of the workflow to run. Practically, only the initial steps of ARPEGGIO, quality check and trimming, can be skipped; otherwise, the workflow will stop for any other step that is set to “False”. Finally, experimental designs are implemented via special modes. By default, ARPEGGIO compares a polyploid species against its two progenitor species (Fig. 1a). With the special mode “POLYPLOID_ONLY”, ARPEGGIO compares a polyploid species from two different experimental conditions (Fig. 1b), while the mode “DIPLOID_ONLY” compares a diploid species from two different conditions (or a polyploid species with an available phased assembly, Fig. 1c).

## Results & Discussion

### Performance of read classification

A simple and common way to analyze polyploid datasets is to concatenate the genome assemblies of the two progenitor species and let the aligner assign a mapping position. The position would define the origin of the read depending on which of the two subgenomes the read was mapped to. We define this approach as the “concatenated” approach.

The performance of EAGLE-RC was assessed using ARPEGGIO while shell scripts were used to evaluate the concatenated approach (see Availability of data and materials). In both cases, the same versions of tools as in ARPEGGIO were used.

For the evaluation, we used six datasets from three pairs of progenitor species that form an allopolyploid or a hybrid, and we compared EAGLE-RC’s classification error to that of the concatenated approach in a similar fashion as (34). In short, each progenitor dataset was treated as an allopolyploid dataset, meaning that all the reads were assigned to a progenitor’s side. With datasets coming from progenitors, the true origin of the reads was known, thus reads assigned to the wrong progenitor’s side were used to calculate a classification error rate.

Two datasets were from *Mimulus guttatus* and *Mimulus luteus*, obtained from (54), with four technical replicates each. Those two species are the progenitors of the allopolyploid *Mimulus peregrinus*. Data from *Gossypium arboreum* and *Gossypium raimondii* was obtained from (29) and consisted of two technical replicates each. Those two species are the progenitors of the hybrid *Gossypium arboreum x raimondii*. The last datasets were produced in-house (Additional File 3) from *Arabidopsis halleri* and *Arabidopsis lyrata* with two biological replicates each. Those two species are the progenitors of the allopolyploid *Arabidopsis kamchatica* (55).

EAGLE-RC showed a lower error rate in all datasets compared to the concatenated approach (Table 1). The error rate was consistently between 3 to 4 times less with EAGLE-RC. When looking at absolute values, the improvement from read classification varied: from changes below 0.1% in *Gossypium* to almost 20% when using *Mimulus* data. These differences could be attributed to many factors, such as divergence between diploids, quality of genome assembly, and sequence data quality. From a qualitative point of view, *Mimulus* had lower quality assemblies compared to the other species, and this difference might also explain the higher error rates in both methods. Also, even though the *Gossypium* data was treated as allopolyploid, the large divergence between the two *Gossypium* species, particularly in terms of genome size, made the read classification task easier (Additional File 4).

**Table 1:** overview of the read assignment accuracy of EAGLE-RC against the concatenation method with real datasets. The first column shows the species name behind the dataset, the second specifies if the replicates consisted of biological or technical replicates, the third and the fourth one show the error rate of the two approaches: concatenating genomes or read classification. The error rate was obtained by the number of reads assigned to the wrong genome divided by the total number of reads that were uniquely mapped and deduplicated.

| Datasets |  | Average number of uniquely mapped reads |  | Average error rate |  |
| --- | --- | --- | --- | --- | --- |
| Species | Type of replicate (#) | Concatenated genome | Read classification | Concatenated genome | Read classification |
| *Arabidopsis halleri* | Biological (2) | 17'258'758 | 18'311'330 | 3.98 % | 1.16 % |
| *Arabidopsis lyrata* | Biological (2) | 22'204'342 | 23'301'056 | 5.94 % | 1.45 % |
| *Mimulus guttatus* | Technical (4) | 1'420'116 | 1'288'800 | 26.78 % | 7.52 % |
| *Mimulus luteus* | Technical (4) | 3'889'458 | 3'760'614 | 9.80 % | 2.29 % |
| *Gossypium arboreum* | Technical (2) | 253'912'667 | 254'261'702 | 0.0044 % | 0.0013 % |
| *Gossypium raimondii* | Technical (2) | 242'590'069 | 246'935'598 | 0.0039 % | 0.0019 % |

Overall, EAGLE-RC showed a lower error rate with minimal loss of reads classified as ambiguous (Additional File 4). On the one hand, EAGLE-RC showed a lower error rate, while on the other, the absolute number of correctly assigned reads was lower in EAGLE-RC compared to the concatenated approach (Additional File 4). This happened because the reads classified as “ambiguous” reduced the amount of the correctly classified reads (both true negative and true positive reads). When focusing on the difference in true positive reads between EAGLE-RC and concatenation, values are negligible for both *Arabidopsis* and *Gossypium* datasets, representing <0.01% of uniquely mapped reads. In the case of *Mimulus*, the number of true positive reads is ~10% higher in the concatenated approach, but the error-rate is also 3 to 4 times higher compared to EAGLE-RC. Taken together, these results suggest that EAGLE-RC has a clear advantage when analyzing allopolyploid WGBS data, where higher accuracy in subgenome recognition is required.

In this evaluation, we have not examined in detail the effect of the genetic divergence between progenitor genomes and allopolyploid genomes. Divergence results from DNA mutations happening after polyploidization and leading to changes on both progenitor sides in the polyploid’s genome. The amount of differences is proportional to the number of generations, i.e. time, since polyploidization. As an example *M. peregrinus* is a 140-years old polyploid, and thus the changes in its genome might be very few. We speculate that ARPEGGIO should be tolerant for older allopolyploids, as both EAGLE-RC and HomeoRoq have shown good performance with both DNA and RNA-seq data of A. kamchatica, which is estimated to have originated around 20,000-250,000 years ago (41,56,57).

### Example run with Mimulus data

To illustrate a full run of ARPEGGIO, we analyzed publicly available data coming from the natural allopolyploid *Mimulus peregrinus* and its progenitors *M. guttatus* and *M. luteus* (58).

First, we downloaded the raw WGBS data consisting of four technical replicates for each species, the genome assemblies of the progenitors with their annotation and a chloroplast genome to check conversion efficiency (details in Availability of data and materials). For WGBS data, genome assemblies and annotations we made sure that all files were formatted according to ARPEGGIO’s user guidelines.

Second, we created a metadata file specifying for each sample the sequencing strategy, single end, and the origin of the samples, i.e. *M. guttatus* samples were labeled “parent1”, *M. luteus* samples “parent2” and *M. peregrinus* samples “allopolyploid”.

With the input files ready, the configuration file was set up in two rounds. In the first round the general parameters were configured with the locations of output folder and metadata file, and data was specified as single end. By default, ARPEGGIO compares allopolyploid to progenitors (Fig 1a), meaning that no specific changes needed to be done to include the experimental design for this dataset. Then, all conditional rules were set to false and ARPEGGIO was run to only perform quality checks. With this round we were able to get more details for the trimming step. In the second round, all the parameters were set for all the different steps in the workflow and all conditional rules were set to true to perform a full run of ARPEGGIO with eight cores. The configuration file, the MultiQC report and ARPEGGIO’s output for the statistical and downstream analyses can be found in Availability of data and materials. The runtime of the full run on a Debian system, using eight CPU cores Intel(R) Xeon(R) CPU E5-4640 at 2.40GHz was approximately 24 hours. The average times for each step can be found in Additional File 2.

After comparing the methylation pattern of *M. peregrinus* to its progenitors, a total of 760 significant DMRs were found in the allopolyploid, most of them coming from the *M. luteus* side (Table 2). Downstream analyses found very few genes overlapping with these significant regions, suggesting that most of the methylation changes occur in intergenic rather than genic regions. For the *M. guttatus* side, 35 genes were found, mostly associated with changes in CG and CHG context, while for the *M. luteus* side only 2 genes were found in CG context. These genes represent a very small proportion of the total number of annotated genes in *M. guttatus*, almost 30’000, and *M. luteus*, almost 50’000. Taken all together, these results suggest almost no change in the global methylation pattern of genes in the natural allopolyploid compared to the two progenitors.

**Table 2:** summary of ARPEGGIO’s downstream analyses on the dataset from Edger and colleagues. The table is divided in two parts, one per progenitor. For each progenitor, the table shows the number of differentially methylated regions (DMRs) for each context, the number of genes overlapping with DMRs and the total number of genes found over all contexts.

|  | *Mimulus guttatus* |  |  | *Mimulus luteus* |  |  |
| --- | --- | --- | --- | --- | --- | --- |
| Methylation context | CG | CHG | CHH | CG | CHG | CHH |
| DMRs | 65 | 126 | 23 | 277 | 211 | 58 |
| Total DMRs | 214 |  |  | 546 |  |  |
| Genes overlapping DMRs | 13 | 20 | 2 | 2 | 0 | 0 |
| Total genes | 35 |  |  | 2 |  |  |

Our analyses use a different approach and different tools compared to (54), but Edger and colleagues also looked at changes in methylation pattern from progenitor to allopolyploid. The authors observed were similar methylation patterns within gene bodies, when comparing progenitors to natural allopolyploids. This is consistent with ARPEGGIO’s downstream analyses showing few genes overlapping with DMRs. Additionally, further analyses in (54) showed that most of the methylation changes happened in transposable elements, another result in agreement with the number of intergenic DMRs found by ARPEGGIO.

### User’s experience and best practices

ARPEGGIO’s user documentation, available through the GitHub Wiki, offers additional information for more and less experienced users. For less experienced users, the documentation offers a step by step guide of how to setup and run ARPEGGIO on a given dataset: data and system requirements, input files needed, configuration file instructions, commands to run the workflow and a map of the output structure. For experienced users, we tried to be as transparent as possible about ARPEGGIO’s code and its architecture to make any customization of scripts and code easier.

As a whole, ARPEGGIO is meant to simplify reproducible data analysis, but best practices, such as data diagnostics and information sharing should be kept in mind. The complete ARPEGGIO pipeline should be run once data quality and potential sources of errors are assessed. To have more control over the analysis process, users also have the option to run ARPEGGIO steps one by one. By modifying the configuration file to add further steps, the workflow will rerun only the parts that need to be updated. To ensure reproducibility when using ARPEGGIO, there are three specifications that need to be included with the datasets: the configuration file settings, the metadata file and the version of ARPEGGIO.

### Software choice

Many alternative tools exist to perform some of ARPEGGIO’s steps. For example, several aligners exist for short-read bisulfite sequencing data such as `bwa-meth` (59), `BSmap` (60), `BitMapperBS` (61), `SNAP` (62) and `gemBS` (63). The `Bismark` suite was selected because it included tools to perform alignment, deduplication and methylation extraction for any context all in one centralized package. Most if not all of the other aligners depend on external packages for downstream analyses of alignment files.

Similarly, many tools exist for DMRs discovery in whole-genome bisulfite sequencing data for all methylation contexts: `BSmooth` (64), `metilene` (65), `MOABS` (66), `BiSeq` (67), `MethylKit` (68) and others (69).

In the case of `dmrseq`, the tool was chosen because of its two step approach: first selecting candidate regions and then evaluating their statistical significance by taking into account both biological variability and spatial correlation. This approach offers important advantages such as limited loss of power and better FDR control, both critical aspects when detecting DMRs (70).

The selection of an appropriate alignment or statistical tool for WGBS data would require an independent benchmark of such tools. An ideal benchmark should evaluate tools on a variety of conditions and provide some guidelines about their suitability and use. Currently, no such benchmarks exist, and a thorough evaluation was out of the scope of this paper. ARPEGGIO provides a convenient implementation of the selected tools and its architecture allows future modifications as long as the input/output structure of the Snakemake rules is preserved. This means that if any of the tools included in the workflow are shown to be underperforming compared to others, ARPEGGIO can be adapted accordingly.

### Comparison to other workflows

To compare ARPEGGIO to other workflows, we selected key steps specifically related to WGBS data analysis (Table 3). The results included workflows able to work with raw bisulfite reads from WGBS and excluded highly specialized (i.e. alignment only or downstream only) and commercial workflows.

**Table 3:**
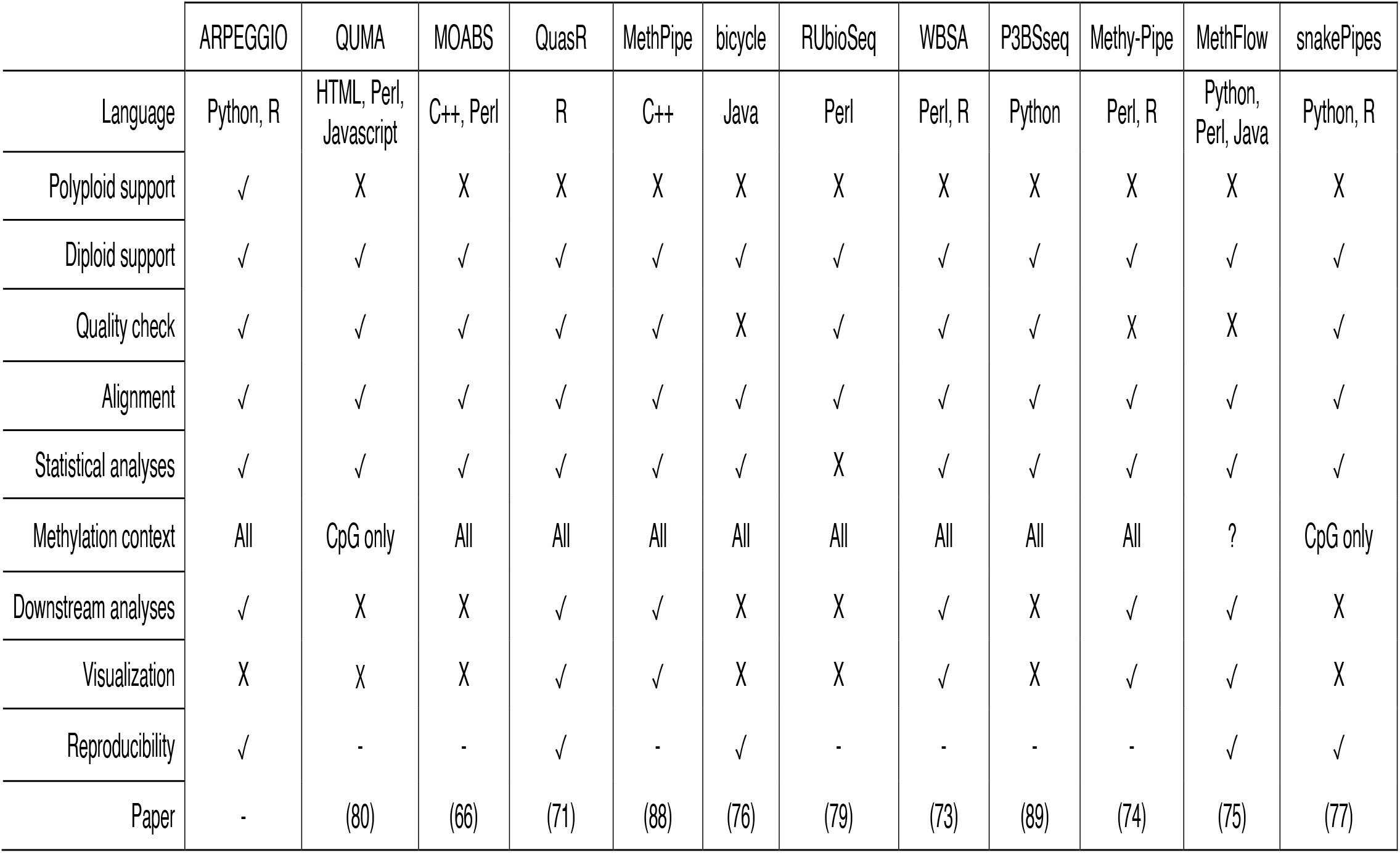
comparison between ARPEGGIO and other available, non-commercial and general workflows able to work with raw WGBS data. There were a total of 12 workflows found and different features were selected for this comparison. The language indicates the main language(s) used to program the workflow. Polyploid support refers to support analysis of data from a polyploid with no official genome assembly available. Diploid support refers to analysis of data from a diploid or a polyploid with an available official genome assembly. Quality check, alignment, statistical and downstream analyses are all different steps in the data analysis process with downstream analyses being defined as follow-up analyses on DMRs found by the statistical analyses. Methylation contexts are 3 in total: CpG, CHG and CHH and this feature is sometimes limited to CpG only. Visualization represents any script or function allowing the user to visualize the DMRs found by the statistical analyses. Reproducibility is difficult to quantify and in this table a tool was considered reproducible if the corresponding paper mentioned reproducibility as one of their goals.

ARPEGGIO is the only workflow specifically targeted at polyploids, making it the main unique feature compared to other available workflows. Other features that were lacking in other workflows, but present in ARPEGGIO, were downstream analyses and reproducibility. Around half of the workflows investigated included downstream analyses (71–75). The lack of this feature might be due to downstream analyses being highly variable according to biological context, question, and aim of the research. With ARPEGGIO, the aim was to consolidate performant tools into a common approach that could be used as a start for further investigation; in our case downstream analyses leading to a list of genes. Reproducibility was another main feature present in ARPEGGIO that was lacking in many workflows, but appeared to be more prevalent in more recent publications (71,75–77). Enhancing and promoting reproducibility is essential to ensure that discoveries stand the test of time (78). Other features were very similar across workflows. All workflows support diploid data, which is considered the same as polyploid data with an available polyploid phased assembly. When comparing the presence of quality check, alignment and statistical analyses, most workflows included them all together, but some didn’t include either quality check (74–76) or statistical analyses (79). For methylation contexts, only two workflows focused on CpG context only (77,80), while all the other allowed analyses for all contexts (CpG, CHG and CHH).

One feature not implemented in ARPEGGIO, but present in other workflows, is visualization of DMRs. This step, similar to downstream analyses, is highly context dependent. The `dmrseq` package offers ways to visualize DMRs, but this was not included in ARPEGGIO. Instead, the workflow outputs an `Rdata` file with all information concerning DMRs that users can use in their custom analyses. It is important to stress that visualization is essential for high-throughput data analysis, and should happen at any step in the data analysis process.

It is important to note that Table 3 focuses only on features related to WGBS data analysis, the only data type supported by ARPEGGIO. Some of the workflows support additional data types and analyses: QuasR supports ChIP-seq, RNA-seq, smRNA-seq and allele-specific data analyses, RUBioSeq supports single-nucleotide and copy number variants (SNVs and CNVs) analyses and snakePipes supports simple DNA-mapping, ChIP-seq, ATAC-seq, HiC, RNA-seq and scRNA-seq data.

Overall, ARPEGGIO was the only workflow supporting polyploid data, and among all the different aspects considered, one of the few workflows including downstream analyses that explicitly set reproducibility as one of its main goals.

## Conclusions

Research on DNA methylation in allopolyploids at a whole genome level seems to be favoring established allopolyploid species (i.e. crops). This can be partially attributed to two factors: 1) challenges in generating allopolyploid genome assemblies; and, 2) a laborious data analysis process. Here we presented ARPEGGIO: the first workflow for the analysis of allopolyploid WGBS data. ARPEGGIO includes a read classification algorithm, EAGLE-RC, to assign allopolyploid reads to the correct progenitor’s side. EAGLE-RC showed better performance against a common concatenation for six different WGBS datasets. Read classification is part of a full set of analyses included in ARPEGGIO, going from raw sequencing data up to a list of genes showing differential methylation. The implementation of ARPEGGIO aimed at ease of use and reproducibility, both essential factors to have an accessible yet up-to-standard tool.

With ARPEGGIO, we provide a first step towards a future of standardized tools and workflows in polyploid research.

## Availability and requirements

Project name: ARPEGGIO

Project home page: https://github.com/supermaxiste/ARPEGGIO

Operating system: Linux

Programming language: Python and R

Other requirements: Python 3, Conda, [Singularity]

License: GNU GPL v3.0

## List of abbreviations

WGBS: Whole genome bisulfite sequencing
DMRs: Differentially methylated regions

## Consent for publication

Not applicable

## Availability of data and materials

Data from cotton taken from (29), available in the NCBI Nucleotide and Sequence Read Archive (SRA) under [SRA:SRP071640]. Data from *Mimulus* taken from (58), available in the NCBI Gene Expression Omnibus (GEO) under [GSE95799]. Data from *Arabidopsis* available in the DDBJ Sequence Read Archive (DRA) under [DRA009902].

The *Gossypium raimondii* v2.0 genome assembly (81) and *Mimulus guttatus* v2.0 (82) genome assembly and annotation were downloaded from Pythozome v12.1 (83). The *Gossypium arboreum* v2_a1 (84) genome assembly was downloaded from CottonGen (85). The *Mimulus luteus* (54) assembly and its annotation were downloaded from Dryad (58). The *Arabidopsis halleri* v2.2 genome assembly was taken from (86) and the *Arabidopsis lyrata* v2.2 genome assembly was taken from (57).

The scripts used for the evaluation of EAGLE-RC and genome concatenation together with the details about the *Mimulus* example run can be found on: https://github.com/supermaxiste/ARPEGGIO_paperAnalyses

## Authors contributions

SM started the bisulfite treatment and library preparation protocol optimization, wrote most of the ARPEGGIO code and manuscript and tested EAGLE-RC. TK updated EAGLE-RC to a new version supporting bisulfite sequencing data and contributed to describe the model with its new features. TK, SD and MR tested ARPEGGIO on their own devices. SD helped with bug fixing in ARPEGGIO, optimized Conda support and implemented Singularity support. RSI managed, coordinated and optimized the bisulfite treatment protocol, library preparation and sequencing process of the *Arabidopsis* samples. LM executed the bisulfite treatment and library preparation. JS, MR, RSI and KS supervised the project and provided important insights at several stages of the project. All authors read and approved the final manuscript.

## Acknowledgements

We thank A. Morishima and M. Wyler for all the support in the DNA extraction, bisulfite treatment and library preparation steps and the optimization of the protocol; the Functional Genomic Center Zurich and M. Hatakeyama for sequencing and data handling; the URPP Evolution in Action program for the opportunity to present ARPEGGIO; the Robinson and Shimizu group members for the feedback during different stages of the project, in particular R. Huang, K. Hembach, S. Orjuela and C. Soneson for the support with Snakemake and the workflow development process.

## Additional file 1

- **Format: pdf**
- **Title of data:** Example of relationships between rules in ARPEGGIO
- **Description:** A graph showing the input/output relationships between different rules in ARPEGGIO in an example “default” run with single end reads.

## Additional file 2

- **Format: pdf**
- **Title of data:** Plot with average runtimes in ARPEGGIO with Mimulus data
- **Description:** A plot with the average runtime for each main step in the ARPEGGIO pipeline: conversion check, quality check, trimming, alignment, deduplication, read classification, methylation extraction and statistical analyses.

## Additional file 3

- **Format: pdf**
- **Title of data:** Plant material and WGBS library synthesis
- **Description:** Details about the plant conditions, sampling, DNA extraction, bisulfite treatment and sequencing strategies.

## Additional file 4

- **Format: pdf**
- **Title of data:** Read statistics about datasets used to compare EAGLE-RC against concatenation method
- **Description:** All the numbers related to the datasets used to compare EAGLE-RC to the concatenation method: total reads, uniquely mapped reads (and not), duplicated reads, correct, ambiguous and wrongly classified reads and error rate.

