## Supplementary figures and images for "ARPEGGIO: Automated Reproducible Polyploid EpiGenetic GuIdance workflOw"

### Additional File 1

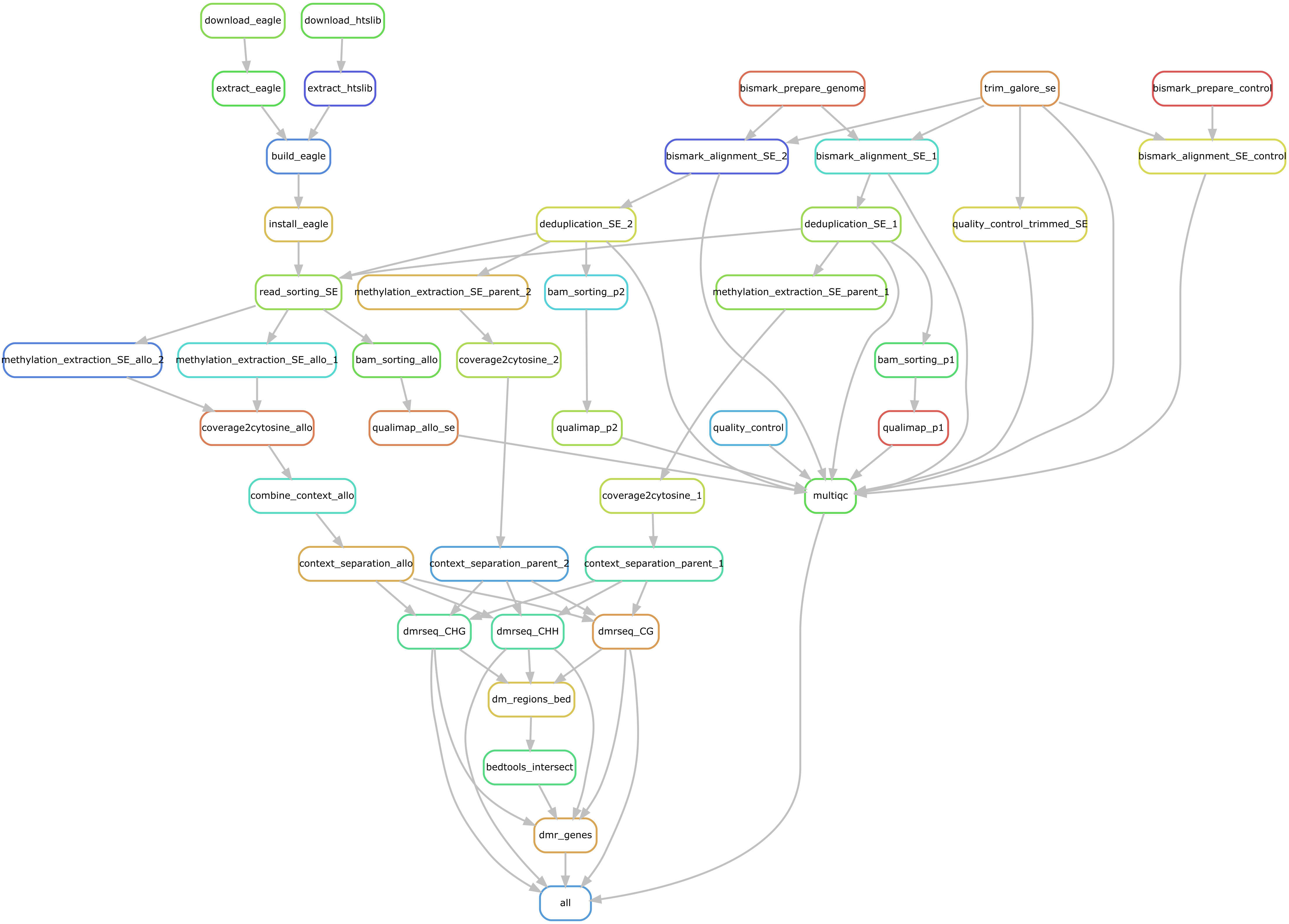

### Additional File 2

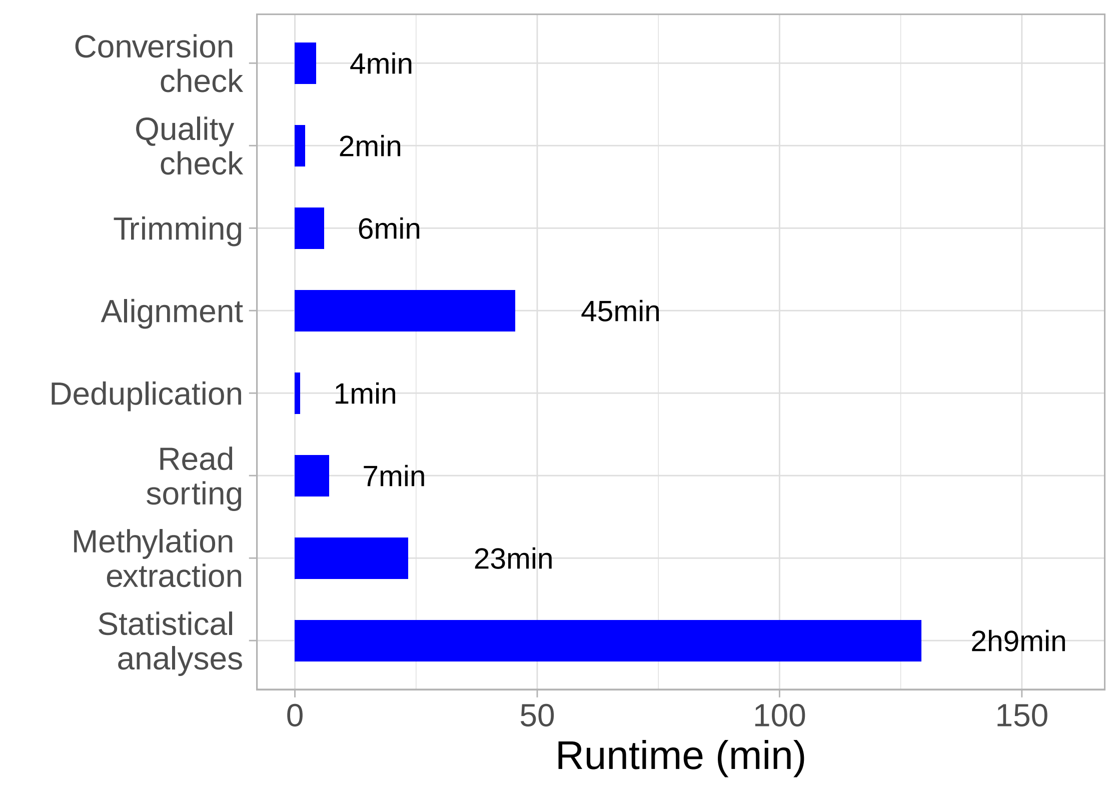
