## Additional File 3 for "ARPEGGIO: Automated Reproducible Polyploid EpiGenetic GuIdance workflOw"

### Plant material and WGBS library synthesis

All plant individuals were incubated in a climate chamber set at 22˚C , with 60% relative humidity, 16 hours light/8hours dark cycles. Mature leaves were collected from each individual, and DNA samples were extracted by DNeasy Plant Mini Kit (Qiagen) from 3 individuals of each species. For the samples *A. halleri* 1 and *A. lyrata* 1, DNA was first treated by bisulfite (BS) using MethylEdge™ Bisulfite Conversion System (Promega), and sequencing library was synthesized by using TruSeq DNA Methylation Kit (Illumina). For the samples *A. halleri* G1 and *A. lyrata* G1, DNA was first synthesized to sequencing library using KAPA HyperPrep Kit with TruSeq DNA Single Indexes Set (Illumina), and BS-treated by EZ DNA Methylation-Gold Kit (ZYMO Research).
The libraries were paired-end sequenced by Illumina HiSeq 4000 (126bp x 2, *A. halleri* 1 and *A. lyrata* 1) and NovaSeq 6000 (150bp x 2, *A. halleri* G1 and *A. lyrata* G1).
