## Additional File 4 for "ARPEGGIO: Automated Reproducible Polyploid EpiGenetic GuIdance workflOw"

**Read statistics about datasets used to compare EAGLE-RC against concatenation method**

The following tables provide all the numbers behind the comparison between EAGLE-RC against the concatenation method. Here's a list with all the samples used and their corresponding accession number:

- *A. halleri*: SAMD00208469
- *A. lyrata*: SAMD00208470
- *A. halleri* G1: SAMD00208471
- *A. lyrata* G1: SAMD00208472
- *M. luteus* 1, 2 , 3 , 4: SRX2618908, SRX2618909, SRX2618910, SRX2618911
- *M. guttatus* 1 , 2 , 3 , 4: SRX2618912, SRX2618913, SRX2618914, SRX2618915
- *G. arboreum* 1 , 2: SRR3219104, SRR3219105
- *G. raimondii* 1, 2: SRR3219088, SRR3219089

| | *A. halleri* | *A. lyrata* | *A. halleri* | *A. lyrata* |
|---|---|---|---|---|
| Method | concatenated | | read sorting | |
| Total reads | 40’160’266 | 44’895’570 | 40’160’266 | 44’895’570 |
| Uniquely mapped | 21’012’844 (52.3%) | 23’067’388 (51.4%) | 22’119’312 (55.3%) | 24’433’418 (54.7%) |
| Not uniquely mapped | 19’147’422 | 21’828’182 | 18’040’954 | 20’462’152 |
| Duplicated reads | 5’293’580 | 7’585’268 | 5’338’990 (24.1%) | 7’897’614 (32.3%) |
| Uniquely mapped and deduplicated | 15’719’264 | 15’482’120 | 16’780’322 | 16’535’804 |
| Correct reads | 15’013’508 (95.5%) | 14’418’656 (91.0%) | 14’953’178 (89.1%) | 14’417’190 (87.2%) |
| Ambiguous | - | - | 1’633’420 | 1’859’612 |
| Wrong reads | 705’756 | 1’063’464 | 193’724 | 259’002 |
| Error % | 4.49 % | 6.87% | 1.30 % | 1.57 % |

| | *A. halleri* G1 | *A. lyrata* G1 | *A. halleri* G1 | *A. lyrata G1* |
|---|---|---|---|---|
| Method | concatenated | | read sorting | |
| Total reads | 80’445’492 | 124’215’824 | 80’445’492 | 124’215’824 |
| Uniquely mapped | 20’321’978 (25.3%) | 31’692’092 (25.5%) | 21’468’920 (26.7%) | 32’969’572 (26.6%) |
| Not uniquely mapped | 60’123’514 | 92’523’732 | 58’976’572 | 91’246’252 |
| Duplicated reads | 1’523’726 | 2’765’528 | 1’626’582 (7.6%) | 2’903’264 (8.8%) |
| Uniquely mapped and deduplicated | 18’798’252 | 28’926’564 | 19’842’338 | 30’066’308 |
| Correct reads | 18’147’370 (96.5%) | 27’480’258 (95.0%) | 18’625’582 (93.9%) | 28’269’980 (94.0%) |
| Ambiguous | - | - | 1’016’786 | 1’399’008 |
| Wrong reads | 650’882 | 1’446’306 | 199’970 | 397’320 |
| Error % | 3.46 % | 5.00% | 1.01 % | 1.32 % |

| | *Mimulus guttatus* 1 | *Mimulus luteus* 1 | *Mimulus guttatus* 1 | *Mimulus luteus* 1 |
|---|---|---|---|---|
| Method | concatenated | | read sorting | |
| Total reads | 11’198’784 | 14’540’898 | 11’198’784 | 14’540’898 |
| Uniquely mapped | 2’612’195 (23.3%) | 6’312’383 (43.4%) | 2’575’518 (23.0%) | 5’908’489 (40.6%) |
| Not uniquely mapped | 8’586’589 | 8’228’515 | 8’623’266 | 8’632’409 |
| Duplicated reads | 1’151’447 (44.1%) | 2’300’413 (36.44%) | 1’249’814 (48.5%) | 2’027’318 (34.3%) |
| Uniquely mapped and deduplicated | 1’460’748 | 4’011’970 | 1’325’704 | 3’881’171 |
| Correct reads | 1’069’658 | 3’623’197 | 976’317 | 3’236’631 |
| Ambiguous | - | - | 250’067 | 556’666 |
| Wrong reads | 391’090 | 388’773 | 99’320 | 87’874 |
| Error % | 26.77% | 9.69% | 7.49% | 2.26% |

| | *Mimulus guttatus* 2 | *Mimulus luteus* 2 | *Mimulus guttatus* 2 | *Mimulus luteus* 2 |
|---|---|---|---|---|
| Method | concatenated | | read sorting | |
| Total reads | 11’125’236 | 14’423’855 | 11’125’236 | 14’423’855 |
| Uniquely mapped | 2’608’418 (23.4%) | 6’291’151 (43.6%) | 2’571’997 (23.1%) | 5’889’892 (40.8%) |
| Not uniquely mapped | 8’516’818 | 8’132’704 | 8’553’239 | 8’533’963 |
| Duplicated reads | 1’149’555 (44.1%) | 2’297’202 (36.5%) | 1’248’440 (48.5%) | 2’026’983 (34.4%) |
| Uniquely mapped and deduplicated | 1’458’863 | 3’993’949 | 1’323’557 | 3’862’909 |
| Correct reads | 1’067’813 | 3’606’521 | 974’850 | 3’219’702 |
| Ambiguous | - | - | 249’601 | 555’449 |
| Wrong reads | 391’050 | 387’428 | 99’106 | 87’758 |
| Error % | 26.81% | 9.70% | 7.49% | 2.27% |

| | *Mimulus guttatus* 3 | *Mimulus luteus* 3 | *Mimulus guttatus* 3 | *Mimulus luteus* 3 |
|---|---|---|---|---|
| Method | concatenated | | read sorting | |
| Total reads | 10’955’877 | 14’136’379 | 10’955’877 | 14’136’379 |
| Uniquely mapped | 2’494’759 (22.8%) | 6’016’962 (42.6%) | 2’459’689 (22.5%) | 5’629’101 (39.8%) |
| Not uniquely mapped | 8’461’118 | 8’119’417 | 8’496’188 | 8’507’278 |
| Duplicated reads | 1’098’495 (44.0%) | 2’193’358 (36.5%) | 1’192’556 (48.5%) | 1’933’785 (34.4%) |
| Uniquely mapped and deduplicated | 1’396’264 | 3’823’604 | 1’267’133 | 3’695’316 |
| Correct reads | 1’022’678 | 3’446’647 | 932’804 | 3’083’952 |
| Ambiguous | - | - | 238’974 | 526’097 |
| Wrong reads | 373’586 | 376’957 | 95’355 | 85’267 |
| Error % | 26.76% | 9.86% | 7.53% | 2.31% |

| | *Mimulus guttatus* 4 | *Mimulus luteus* 4 | *Mimulus guttatus* 4 | *Mimulus luteus* 4 |
|---|---|---|---|---|
| Method | concatenated | | read sorting | |
| Total reads | 10’646’892 | 13’738’854 | 10’646’892 | 13’738’854 |
| Uniquely mapped | 2’433’041 (22.9%) | 5’866’025 (42.7%) | 2’398’688 (22.5%) | 5’485’383 (39.9%) |
| Not uniquely mapped | 8’213’851 | 7’872’829 | 8’247’915 | 8’253’471 |
| Duplicated reads | 1’068’453 (43.9%) | 2’137’718 (36.4%) | 1’159’882 (48.4%) | 1’882’322 (34.3%) |
| Uniquely mapped and deduplicated | 1’364’588 | 3’728’307 | 1’238’806 | 3’603’061 |
| Correct reads | 998’982 | 3’357’807 | 911’356 | 3’003’804 |
| Ambiguous | - | - | 233’955 | 515’291 |
| Wrong reads | 365’606 | 370’500 | 93’495 | 83’966 |
| Error % | 26.79% | 9.94% | 7.55% | 2.33% |

| | *Gossypium arboreum* 1 | *Gossypium raimondii* 1 | *Gossypium arboreum* 1 | *Gossypium raimondii* 1 |
|---|---|---|---|---|
| Method | concatenated | | read sorting | |
| Total reads | 432’844’852 | 356’699’260 | 432’844’852 | 356’699’260 |
| Uniquely mapped | 279’996’748 (64.7%) | 273’814’708 (76.8%) | 280’386’284 (64.8%) | 278’732’236 (78.1%) |
| Not uniquely mapped | 152’848’104 | 82’884’552 | 152’458’568 | 77’967’024 |
| Duplicated reads | 18’846’046 (6.7%) | 13’284’372 (4.8%) | 18’879’306 (6.7%) | 13’697’054 (4.9%) |
| Uniquely mapped and deduplicated | 261’150’702 | 260’530’336 | 261’506’978 | 265’035’182 |
| Correct reads | 260’132’278 | 259’423’814 | 260’022’724 | 262’945’068 |
| Ambiguous | - | | 1’132’490 | 1’613’104 |
| Wrong reads | 1’018’424 | 1’106’522 | 351’764 | 477’010 |
| Error % | 0.00390% | 0.00425% | 0.00134% | 0.00179% |

| | *Gossypium arboreum* 2 | *Gossypium raimondii* 2 | *Gossypium arboreum* 2 | *Gossypium raimondii* 2 |
|---|---|---|---|---|
| Method | concatenated | | read sorting | |
| Total reads | 414’743’906 | 299’026’128 | 414’743’906 | 299’026’128 |
| Uniquely mapped | 264’247’068 (63.7%) | 235’500’124 | 264’620’650 (63.8%) | 240’038’096 (80.3%) |
| Not uniquely mapped | 150’496’838 | 63’526’004 | 150’123’256 | 58’988’032 |
| Duplicated reads | 17’572’436 | 10’850’322 (4.6%) | 17’604’224 | 11’202’082 (4.7%) |
| Uniquely mapped and deduplicated | 246’674’632 | 224’649’802 | 247’016’426 | 228’836’014 |
| Correct reads | 245’718’300 | 223’623’540 | 245’626’594 | 226’937’338 |
| Ambiguous | - | - | 1’059’410 | 1’452’404 |
| Wrong reads | 956’332 | 1’026’262 | 330’422 | 446’272 |
| Error % | 0.00389% | 0.00457% | 0.00134% | 0.00195% |
